# Single Cell T Cell Receptor Repertoire Profiling for Dogs

**DOI:** 10.1101/2021.06.29.450365

**Authors:** Zachary L. Skidmore, Hans Rindt, Shirley Chu, Bryan Fisk, Catrina Fronick, Robert Fulton, Mingyi Zhou, Nathan J. Bivens, Carol N. Reinero, Malachi Griffith, Jeffrey N. Bryan, Obi L. Griffith

**Author notes:** corresponding authors Address correspondence to: Jeffrey N. Bryan,; Obi L. Griffith. These authors contributed equally (Zachary L. Skidmore, Hans Rindt).

## Abstract

**Background:** Spontaneous cancers in companion dogs are increasingly recognized as robust models of human disease. This recognition has led to translational clinical trials in companion dogs with osteosarcoma, lymphoma, melanoma, squamous cell carcinoma, and soft tissue sarcoma. The ability to precisely track tumor-specific immune responses in such clinical trials would benefit from reagents to perform species-specific single cell T cell receptor sequencing (scTCRseq). This technology defines clones of T cells reacting to immune interventions and can help identify the specific epitope of response. Single cell gene expression data give insights into the activity and polarization of the T cell. To date, scTCRseq has not been demonstrated for canine samples.

**Methods:** Samples from two responding dogs in a trial of an autologous deglycosylated melanoma vaccine were selected to demonstrate applicability of scTCRseq in a cancer immunotherapy setting. A single-cell suspension of cryopreserved peripheral blood mononuclear cells (PBMC) was prepared for 10X single cell sequencing. Full length 10X cDNA was amplified using a custom-designed nested PCR of the alpha/beta V(D)J region. A library made from this enriched product (scTCRseq) and a 10X gene expression (GEX) library (scRNAseq) were sequenced on the NovaSeq 6000.

**Results:** 1,850-2,172 estimated V(D)J-expressing cells yielded 87-103.7 million reads with 73.8%-75.8% mapped to a V(D)J gene (beta/alpha chains ratio 1.5:1). 43 TRAJ, 29 TRAV, 12 TRBJ, and 22 TRBV gene segments were observed representing 72.9%, 51.8%, 100%, and 62.9% of all known V and J gene segments respectively. A large diversity of clonotypes was captured with 966-1,253 TRA/TRB clonotypes identified. Both dogs also exhibited a small number of highly abundant T cell clonotypes suggesting the presence of an anti-tumor T cell population. GEX enriched libraries successfully defined large clusters of CD8+ and CD4+ T cells that overlapped with V(D)J-expressing cells.

**Discussion:** The developed reagents successfully generated scTCRseq data, for the first time, which allowed the T cell repertoire to be surveyed in dogs responding to anti-tumor immunotherapy. These reagents will allow longitudinal tracking of anti-tumor T cell dynamics in canine cancer immunotherapy trials.

## Introduction

The biomedical research community, as well as the National Cancer Institute, have recognized the value of companion dog spontaneous models of cancer with release of several targeted funding programs. This recognition has led to the completion of translational clinical trials in companion dogs studying osteosarcoma^1^, lymphoma^2^, melanoma, squamous cell carcinoma, and soft tissue sarcoma^3^. Recently, immunotherapy trials evaluated a human chimeric HER2 Listeria vaccine in osteosarcoma as well as an antibody-linked IL-12 conjugate in melanoma of companion dogs^4^. The outbred genetic variability, body size, cancer-conditioned immune system, and shared environment of companion dogs and humans have been cited as particular advantages of dogs as pre-clinical or interposed post-clinical subjects in veterinary clinical trials of interventions destined for human application^5^.

Immunotherapy has become increasingly important in the treatment of cancer with the rise of checkpoint blockade inhibitors, personalized cancer vaccines, CAR T-cell therapies, and more. As a result, profiling the T cell receptor (TCR) repertoire has become an important tool for understanding, measuring, tracking and even predicting T cell mediated immune responses in a cancer setting^6^. The ability to accurately profile the TCR repertoire can also provide a powerful means of diagnosing and tracking T cell malignancies (e.g., minimal residual disease monitoring). Finally, TCR profiling may play an important role in understanding and treating a host of other immune-related (e.g., autoimmune) diseases.

It has also become increasingly apparent in recent literature that the tumor microenvironment (TME) plays a broad and complex role in tumorigenesis and response to many cancer therapies^7^. In particular the TME is known to be capable of producing an immunosuppressive effect. For example the presence of cytokines such as IL-6, IL-12, and TGFβ can increase PD-L1 expression allowing immune escape of tumor cells via checkpoint pathways^8^. The ability to examine the T cell repertoire, in the context of the TME, using single cell approaches, in a model with a functionally similar environment to that of humans could yield significant insights into the role of the TME in success or failure of immune therapy. Furthermore, single cell TCR profiling facilitates the reconstruction of the complete TCR by matching the alpha-beta pairs of the TCR. This in turn may allow for refinement of TCR-Neoantigen-MHC binding prediction algorithms. Such refinement could have implications for personalized therapy, allowing researchers to determine if a T-cell is reactive to a given neoantigen and opening avenues into engineering T-cells matched to an individual tumor profile.

A limitation of the companion dog model of cancer is the relative paucity of molecular profiling reagents available compared to human or mouse settings. Dog genome-specific reagents are required to effectively evaluate tumor-specific immune responses in cancer immunotherapy trials. While studies have piloted scRNAseq for canine samples^9,10^, to our knowledge comprehensive single cell profiling of the TCR repertoire (scTCRseq) of canine samples has not been demonstrated. There are no published protocols or commercially available kits for popular single cell platforms like the Chromium Single Cell TCR Amplification kits available for human and mouse cells. Here we address this gap by developing and validating canine alpha (TRA) and beta (TRB) chain TCR single cell profiling for the Chromium 10X platform. We show robust isolation of individual canine cells, including a large proportion of T cells, and establish the first detailed single cell TCR repertoires for dog cells.

## Methods

### Samples and single cell suspension

To demonstrate applicability of canine TCR profiling in a cancer immunotherapy setting we identified two individual dogs (Dog_A and Dog_B) with metastatic melanoma from an ongoing immune therapy trial (See **Suppl Table 1** for complete clinical information). Both dogs were treated with autologous deglycosylated vaccines derived from primary and metastatic tumor cultures and showed evidence of clinical response. Dog_A had a progression free interval of 280 days after the first dose of the autologous vaccine. Dog_B had resolution of progressive pulmonary nodules following the vaccine series and survived more than a year without clinical recurrence. Ficoll-separated peripheral blood mononuclear cells (PBMCs) were obtained from each dog and cryopreserved. PBMCs for Dog_A were collected at the time of the first tyrosinase vaccine (Oncept) and for Dog_B were collected 14 days after the initial autologous vaccine treatment. Dog_B was free of macroscopic disease at the time of blood draw. Cryopreserved PBMCs were thawed and a single cell suspension of approximately 20,000 cells was generated according to the 10x Genomics Demonstrated Protocol (CG00039 Rev D). Viability was greater than 90% as assessed by trypan blue exclusion staining.

### Primer design

Primer design and application (**Figure 1**) was modeled from the Chromium Single Cell V(D)J Reagent Kits User Guide (CG000086 Rev L). This protocol uses a nested PCR design. The two forward primers from the 10x protocol were left unchanged. In the first cycle, the forward primer (TRA/TRB Forward 1) primes at the Illumina read 1 (R1) sequencing adapter sequence that is incorporated during generation of 10x Barcoded, full-length cDNA from polyadenylated mRNA. This primer includes a 5’ tail sequence consisting of the P5 priming site used in Illumina sequencing. In the nested primer set, the forward primer (TRA/TRB Forward 2) primes at the P5 sequence and a portion of R1. The forward primers correspond to the 5’ end of 10x cDNA fragments. In the case of full length TCR cDNA this would represent the V gene segment end of the cDNA fragment. Neither forward primer has any specificity for the TCR which comes entirely from the reverse primers. The first reverse primer (TRA/TRB Reverse 1, Outer) primes at the constant (C) region gene segment. The second reverse primer (TRA/TRB Reverse 2, Inner) similarly primes at the C region at an inner, 5’ position relative to the outer primer. In order to design appropriate 3’ primers for use with canine cells we first constructed (*in silico*) a reference complete V(D)JC TCR cDNA sequence along with 10X adapter sequences for both alpha (TRA) and beta (TRB) chains. The alpha chain was based on a representative dog TCR alpha rearranged partial mRNA (GenBank: M97511.1). The closest V gene segment to this partial mRNA was determined by blast alignment against canine V gene sequences in the IMGT database. The constructed cDNA was extended to include (from 3’ to 5’) the Illumina R1 adaptor, 16 nucleotide (nt; 16 x N) 10x cell barcode, 10 nt (10 x N) unique molecular identifier (UMI), 13 nt template switch oligo (TSO), complete V gene segment, complete J gene segment, and complete C gene segment. The beta chain was based on a representative dog TCR beta rearranged partial mRNA (GenBank: HE653957.1). The closest V and C gene segments to this partial mRNA were determined by blast alignment against canine V and C gene sequences in the IMGT database. The constructed cDNA was extended to include (from 3’ to 5’) the R1 adaptor, 10x cell barcode, UMI, TSO, V, D, J, and C gene segments as described above. These constructed cDNA sequences were then used as input to primer3plus (4.0)^11,12^, with forward primers provided as described above, and a target region for reverse primer design specified in the C region. The product of the first (outer) design was used as input for the second (inner) design. Primer oligonucleotides were ordered from Integrated DNA Technologies (Coralville, IA). Final primer sequences are included in **Table 1**.

**Figure 1.**
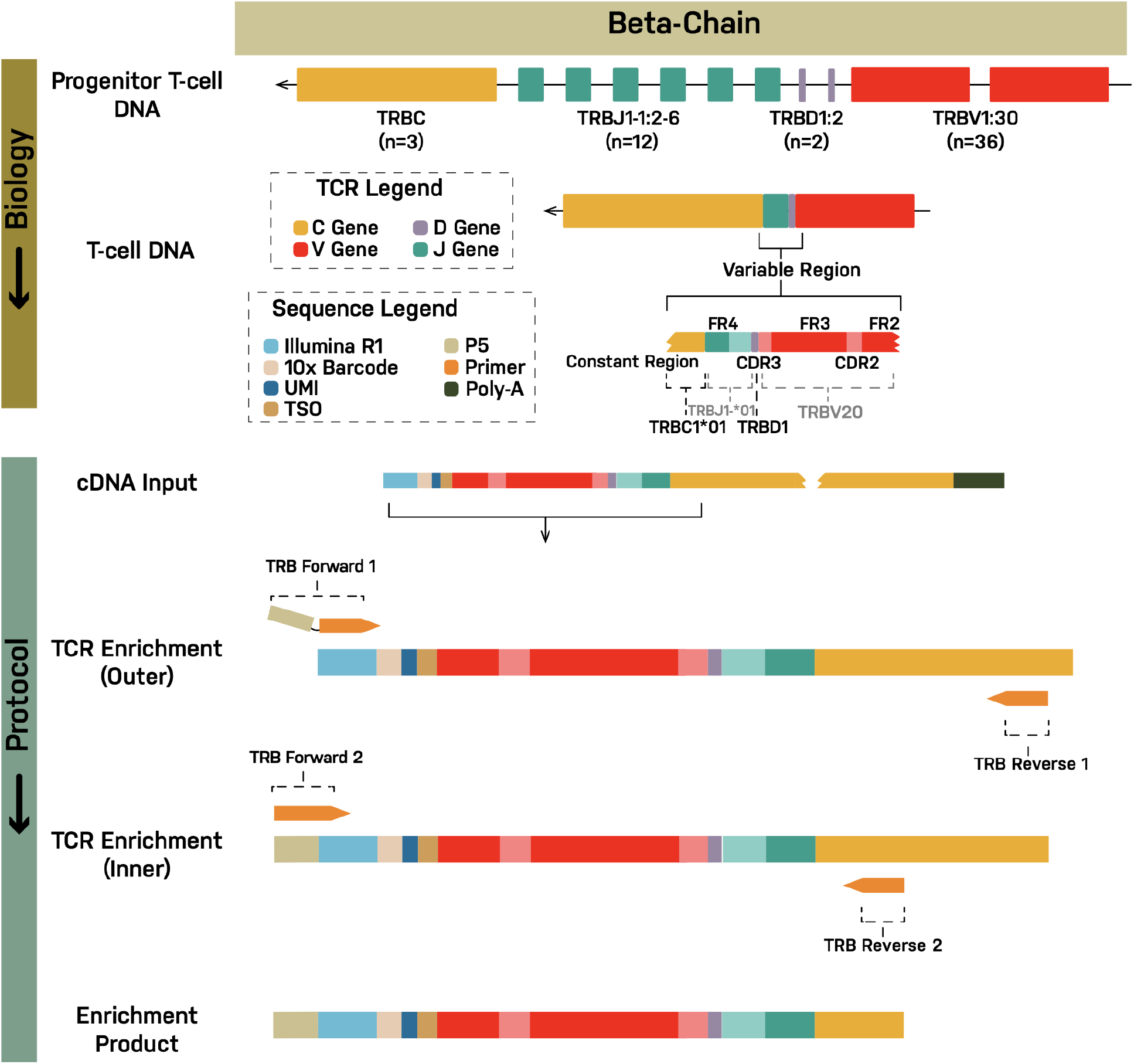
TCR enrichment strategy for use with 10X scRNA sequencing. The TCR V(D)J enrichment strategy is depicted using the beta chain for illustration (See **Suppl Figure 1** for alpha chain). At top, the genomic un-rearranged TRB locus is shown (Note: TRB is located on the negative strand in dogs). During T cell development from progenitor T cells to mature T cells, individual V, D, J and C gene segments are rearranged by somatic recombination to produce a functional TRB locus. Transcription and splicing produce a pre-mRNA and then mRNA for the complete V(D)JC transcript sequence (not shown). In the modified 10X protocol, mRNA (including TRB mRNA) is converted to cDNA. TRB cDNA is then amplified using a nested PCR design. The two forward primers from the 10x protocol were left unchanged. In the first cycle, the forward primer (TRB Forward 1) primes off the Illumina read 1 (R1) sequencing adapter that is incorporated during generation of cDNA. TRB Forward 1 includes a tail sequence consisting of the P5 priming site used in Illumina sequencing. In the second cycle, the forward primer (TRB Forward 2) primes off the P5 sequence and a portion of R1. The first reverse primer (TRB Reverse 1, Outer) primes off the constant (C) region gene segment. The second reverse primer (TRB Reverse 2, Inner) similarly primes off the C region at an inner, 5’ position relative to the outer primer. The beta chain primer design was based off of a dog TCR beta rearranged partial mRNA (GenBank: HE653957.1) which was extended to include (from 3’ to 5’) the R1 adaptor, 10x cell barcode, UMI, TSO, V, D, J, and C gene segments. The constructed cDNA sequence was then used as input to primer3plus (4.0), with forward primers provided as described above, and a target region for reverse primer specified in the C region. The product of the first (outer) design was used as input for the second (inner) design.

**Table 1.**
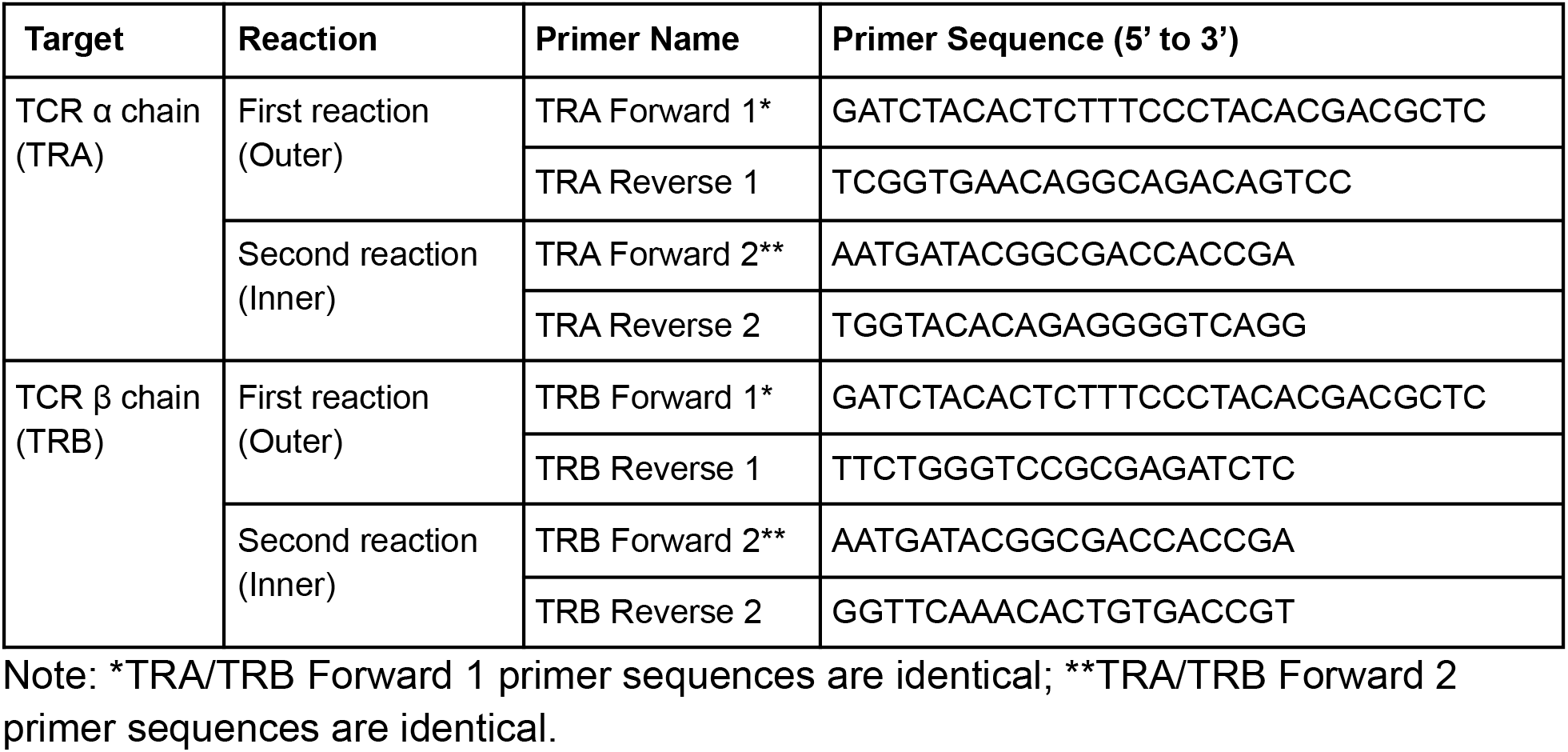
TCR amplification primer sequences.

### cDNA generation

cDNA generation was performed according to the Chromium Single Cell V(D)J Reagent Kits User Guide (CG000086 Rev L), with the exception of the TCR amplification (described below). Briefly, approximately 20,000 cells suspended in phosphate-buffered saline containing 0.4% bovine serum albumin were used for Gel Bead-in-Emulsion (GEM) generation, barcoding, post-GEM clean-up, and cDNA amplification. Quality controls were performed using the Agilent Bioanalyzer high sensitivity chip. 2 uL of cDNA were used to separately amplify TCR α and β chains using custom dog-specific primers (see Primer design; TCR amplification). The remaining cDNA were set-aside for gene expression (GEX) profiling.

### TCR amplification

In order to amplify dog TCR sequences, the nested PCR approach utilized for human and mouse protocols was adopted, with dog-specific reverse primers located in the constant region of the TCR cDNA (see Primer design; **Table 1**). 2 uL of cDNA were amplified in a Mastercycler Gradient instrument (Eppendorf) in 100 uL total reaction volume using Phusion high fidelity polymerase (Thermo Fisher). The first reaction consisted of an initial denaturation step (98 °C for 45 s) followed by 12 cycles of 98 °C for 20 s, 65 °C for 30 s, and 72 °C for 60 s. An additional extension step (72 °C for 60 s) was added at the end. 10 uL of the resulting product was amplified in the second reaction. Amplification conditions were identical except for the annealing temperature which was 62 °C. The amplification product was purified using SPRI beads before quality control on the Agilent Bioanalyzer high sensitivity chip and further processing for library generation.

### Library generation

TCR enriched PCR products and set-aside GEX cDNAs underwent library construction and final quality control as described in Chromium Next GEM Single Cell 5’ Reagent Kits v2 (Dual Index) User Guide (CG000331 Rev A).

### Sequencing

GEX and TCR VDJ enriched libraries, produced by the MU Core (as described above), were shipped to the McDonnell Genome Institute for sequencing. The concentration of each 10x single cell and V(D)J library was determined through qPCR (Kapa Biosystems) to produce cluster counts appropriate for the NovaSeq 6000 platform. 500M read pairs were targeted for each gene expression library and 25M read pairs were targeted for each V(D)J library. The libraries were sequenced on the S4 300 cycle kit flow (2×151 paired end reads) using the XP workflow as outlined by Illumina.

### Data processing

All data processing and subsequent analysis made use of Cell Ranger 5.0.1 unless otherwise noted. Raw fastq files were generated from Illumina instrument data using *cellranger mkfastq*. A canine-specific reference package was created using *cellranger mkref* with CanFam3.1 (INSDC Assembly GCA_000002285.2, Sep 2011) canine reference genome and *cellranger mkgtf* filtered CanFam3.1 GTF from Ensembl v102 as input. The GTF file was filtered according to the example provided in the documentation on the 10X Genomic site^13^. A canine-specific V(D)J reference package was created using the *fetch-imgt* script and *cellranger mkvdjref*. The *fetch-imgt* script failed to obtain C-REGION sequences causing errors in subsequent steps. These sequences (IMGT000004|TRAC*01, IMGT000005|TRBC1*01, HE653929|TRBC1*02, and HE653929|TRBC2*01) were obtained from IMGT^14^, artificially spliced, and added manually to the *fetch-imgt* file before completing V(D)J reference generation (**Suppl Data 1**). Single cell feature counts were generated with *cellranger count*. Counts were combined, normalized, and batch corrected across multiple libraries with *cellranger aggregate*. Single cell VDJ sequence assembly and paired clonotype calling was performed with *cellranger vdj*. Genes present in less than 10 cells and cells with less than 100 genes were filtered out of the dataset. Cells whose mitochondrial expression was in the top 5% across all cells were filtered out and DoubletFinder^15^ was used to identify and filter out expected doublets. DoubletFinder was run with parameters derived using the example provided at https://github.com/chris-mcginnis-ucsf/DoubletFinder. Cells were annotated to cell types with SingleR^16^ using expression profiles derived from the DMAP dataset available on Haemopedia, a hematopoiesis cell expression database^17^. The DMAP dataset was converted from Affymetrix probes to human gene IDs and then to dog gene names using the probe to human gene name mapping available on Haemopedia and the human to dog gene mapping acquired from Ensembl v102 BioMart. If different probes mapped to the same human gene the probe with the highest coefficient of variance across all samples in the dataset was kept and all other probes that mapped to that gene were discarded. If a probe entry mapped to multiple genes then that probe-to-gene mapping was duplicated for each gene it mapped to. For the human gene ID to dog gene name conversion all many-to-many mappings were removed, mappings that had multiple human genes to a single dog gene were removed, and all the mappings from one human gene to multiple dog genes were duplicated for each dog gene. Cell types were manually simplified to B cell (B), early B-cell (PreB), basophil (BASO), CD4+ T cell (CD4), CD8+ T cell (CD8), common myeloid progenitor (CMP), dendritic cell (DC), eosinophil (EOS), erythroid (ERY), granulocyte/monocyte progenitor (GMP), granulocyte (GRAN), hematopoietic stem cell (HSC), megakaryocyte (MEGA), megakaryocyte/erythroid progenitor (MEP), monocyte (MONO), and NK cell (NK). All additional ad hoc analysis and figure generation was performed in the Loupe Browser (version 5.0.1), Loupe VDJ Browser (version 4.0.0) and Seurat^18^ (version 4.0.1).

## Results

### Single cell V(D)J enrichment and V(D)J/GEX sequencing results

Nested PCR primers were successfully designed for dog TCR alpha (TRA) and TCR beta (TRB) chains (**Figure 1**; **Suppl Figure 1; Table 1**). PCR products of the expected size (approximately 650 bp) were observed by gel electrophoresis (**Figure 2**). Sequencing libraries from TCR enriched products (TRA and TRB) and unenriched GEX cDNA were successfully generated, for both dogs, with suitable quantities, that also showed expected fragment size distributions (**Figure 3; Suppl Figure 2**). Sequencing data metrics approximated comparable human data for the TCR enriched libraries (**Table 2**)^19,20^. We observed 1,850-2,172 estimated V(D)J expressing cells with mean reads per cell of 47,042-47,745 and fraction of reads in cells of 39.8%-48.0%. In total, 87.0-103.7 million reads were generated for the two V(D)J libraries with 96.3-96.6% valid barcodes. Q30 bases for barcodes, Read 1, Read 2, and UMI sequences were greater than 89.8% in all cases. For the GEX enriched libraries we also obtained results comparable to human data (**Table 3**)^21^. We observed 6,968-7,448 estimated cells with mean reads per cell of 69,066-72,254 and fraction of reads in cells of 95.9-96.1%. In total, 503.5-514.4 million reads were generated for the two GEX libraries with 92.8-93.4% valid barcodes. Q30 bases for barcodes, Read 1, Read 2, and UMI sequences were greater than 84.5% in all cases. For gene estimation purposes, 73.0%-73.4% of total reads mapped to the genome with 14,610-14,751 genes detected and 1,532-1,635 median genes per cell.

**Table 2.** Single cell V(D)J sequencing metrics.

|  | **Metric** | **Dog_A** | **Dog_B** |
| --- | --- | --- | --- |
| **Cells** | Estimated number of cells | 1,850 | 2,172 |
|  | Mean reads/cell | 47,042 | 47,745 |
|  | Mean used reads/cell | 16,322 | 19,839 |
|  | Number Cells with Productive V-J Spanning Pair | 846 (45.7%) | 758 (34.9%) |
|  | Fraction reads in cells | 39.8% | 48.0% |
| **Enrichment** | Reads Mapped to Any V(D)J Gene | 73.8% | 75.8% |
|  | Reads Mapped to TRA | 31.2% | 30.0% |
|  | Reads Mapped to TRB | 42.4% | 45.5% |
| **V(D)J Expression** | Median TRA UMIs per Cell | 6.0 | 4.0 |
|  | Median TRB UMIs per Cell | 12.0 | 15.0 |
| **V(D)J Annotation** | Cells With Productive V-J Spanning Pair | 45.7% | 34.9% |
|  | Paired Clonotype Diversity | 83.3 | 19.4 |
|  | Cells With TRA Contig | 70.6% | 53.3% |
|  | Cells With TRB Contig | 91.2% | 94.3% |
|  | Cells With CDR3-annotated TRA Contig | 66.3% | 48.8% |
|  | Cells With CDR3-annotated TRB Contig | 87.4% | 92.1% |
|  | Cells With V-J Spanning TRA Contig | 68.5% | 50.9% |
|  | Cells With V-J Spanning TRB Contig | 87.9% | 92.3% |
|  | Cells With Productive TRA Contig | 62.3% | 45.2% |
|  | Cells With Productive TRB Contig | 83.5% | 89.7% |
| **Sequencing** | Number of Reads | 87,026,777 | 103,702,506 |
|  | Valid Barcodes | 96.3% | 96.6% |
|  | Q30 Bases in Barcode | 96.8% | 96.7% |
|  | Q30 Bases in RNA Read 1 | 93.8% | 93.8% |
|  | Q30 Bases in RNA Read 2 | 89.8% | 90.0% |
|  | Q30 Bases in UMI | 96.2% | 96.2% |

Note: Paired Clonotype Diversity is computed as the Inverse Simpson Index of the clonotype frequencies. A value of 1 indicates a minimally diverse sample - only one distinct clonotype was detected. A value equal to the estimated number of cells indicates a maximally diverse sample.

**Table 3.** Single cell gene expression (GEX) sequencing metrics.

|  | **Metric** | **Dog_A** | **Dog_B** |
| --- | --- | --- | --- |
| **Cells** | Estimated number of cells | 6,968 | 7,448 |
|  | Fraction Reads in Cells | 95.9% | 96.1% |
|  | Mean Reads per Cell | 72,254 | 69,066 |
|  | Median Genes per Cell | 1,532 | 1,635 |
|  | Total Genes Detected | 14,610 | 14,751 |
|  | Median UMI Counts per Cell | 4,422 | 4,801 |
| **Sequencing** | Number of Reads | 503,465,164 | 514,405,307 |
|  | Number of Short Reads Skipped | 0 | 0 |
|  | Valid Barcodes | 92.8% | 93.4% |
|  | Valid UMIs | 99.8% | 99.9% |
|  | Sequencing Saturation | 80.1% | 77.4% |
|  | Q30 Bases in Barcode | 87.9% | 88.4% |
|  | Q30 Bases in RNA Read 1 | 84.5% | 85.1% |
|  | Q30 Bases in RNA Read 2 | 85.8% | 84.9% |
|  | Q30 Bases in UMI | 87.8% | 88.2% |
| **Mapping** | Reads Mapped to Genome | 73.0% | 73.4% |
|  | Reads Mapped Confidently to Genome | 70.1% | 71.0% |
|  | Reads Mapped Confidently to Intergenic Regions | 7.5% | 8.2% |
|  | Reads Mapped Confidently to Intronic Regions | 6.5% | 7.2% |
|  | Reads Mapped Confidently to Exonic Regions | 56.1% | 55.6% |
|  | Reads Mapped Confidently to Transcriptome | 48.0% | 47.2% |
|  | Reads Mapped Antisense to Gene | 2.5% | 2.6% |

**Figure 2.**
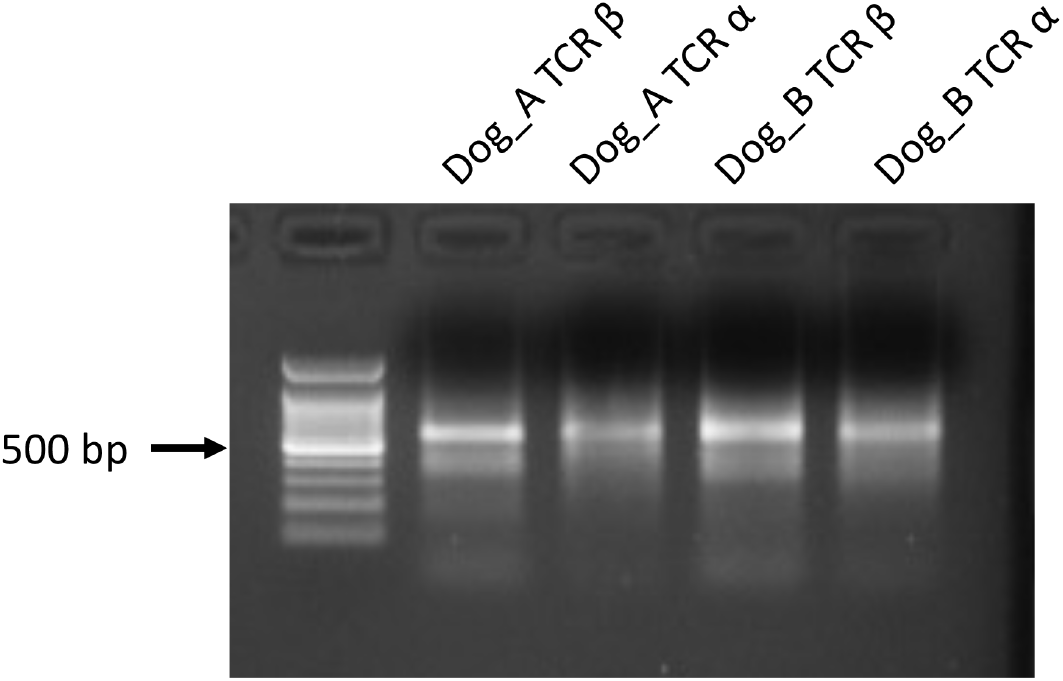
Gel image of final TCR amplification product (Dog_A and Dog_B). Gel electrophoresis result visualizing product of TCR-alpha (TRA) and TCR-beta (TRB) nested PCR amplification for two dogs using custom primers (**Table 1**) and a modified Chromium 10X protocol (Methods). Gel bands of the expected size (∼650bp) were observed for all 4 reactions.

**Figure 3.**
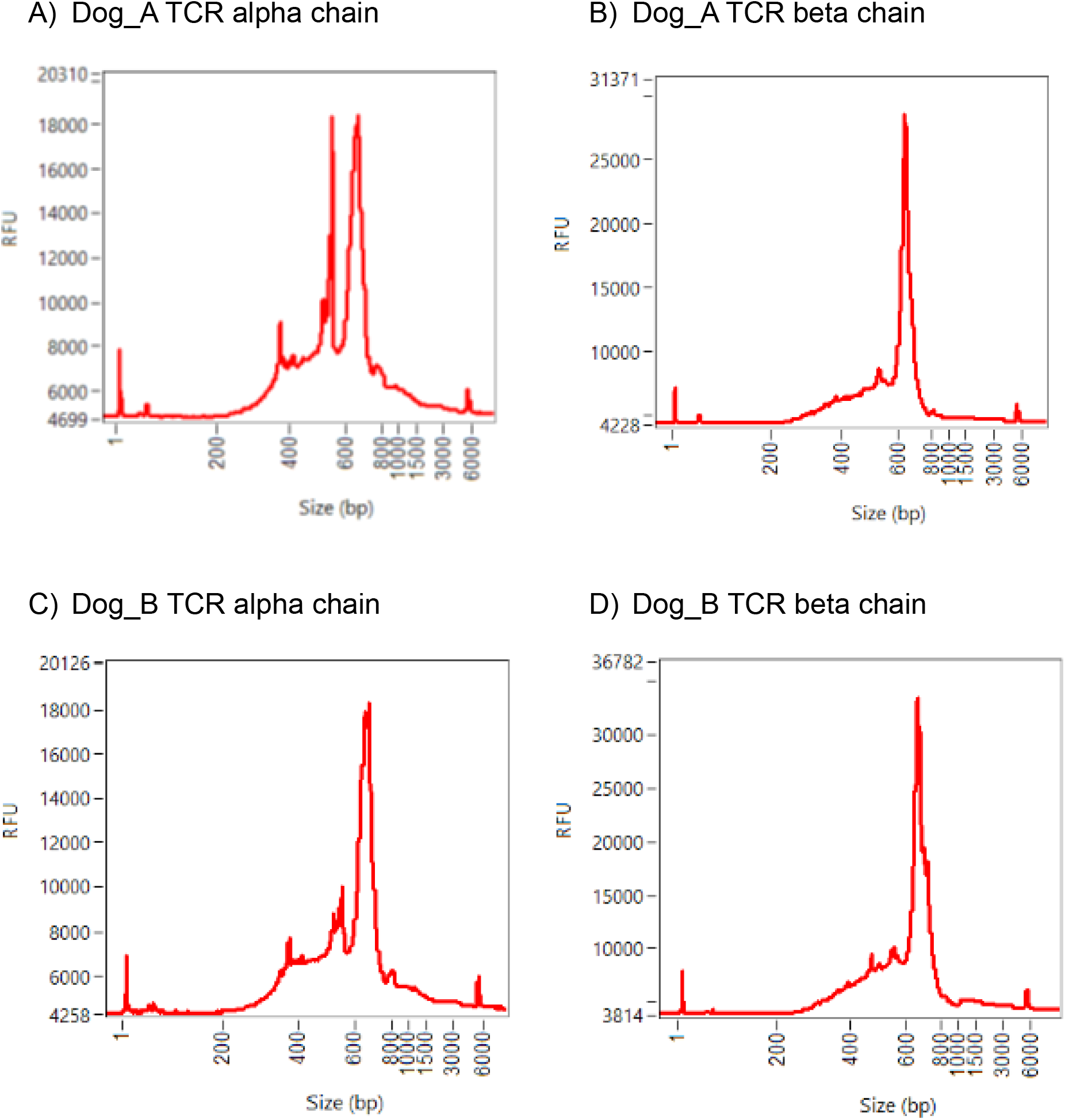
Analytic traces of the libraries prepared from the four TCR amplification products by the fragment analyzer. Sequencing libraries were constructed from the 4 purified TCR amplification products and analyzed on the Agilent Fragment Analyzer using the NGS Fragment kit (DNF-473-500). The trace profiles are unique because they are derived from smaller amplified PCR products.

### Single cell V(D)J repertoire

For TCR enriched libraries 73.8%-75.8% of reads mapped to a V(D)J gene with an approximate ratio between beta/alpha chains of 1.5:1. Between 45.2-62.3% of cells exhibited a productive TRA contig. Similarly between 83.5-89.7% of cells exhibited a productive TRB contig. Cells with a productive V-J spanning pair ranged from 34.9-45.7% (**Table 2**). We observed 43 TRAJ, 29 TRAV, 12 TRBJ, and 22 TRBV unique gene segments at some level of support. These observations represent 72.9%, 51.8%, 100%, and 62.9% of all known V and J gene segments respectively **(Figure 4; Suppl Figure 3)**. Both Dog_A and Dog_B exhibited relatively clonal TCR repertoires, with paired clonotype diversity (Inverse Simpson Index) equal to 83.3 and 19.4 respectively (**Figure 5, Suppl Figure 4, Table 2**). Each dog had one or a few CDR3 clonotypes accounting for a substantial fraction of all cell barcodes (**Suppl Table 2-3**). For example, for Dog_B, just two TRA and two TRB clonotypes account for approximately 20% and 40% of all barcodes respectively (**Figure 5**). At the same time, a large diversity of clonotypes was captured with 966-1,253 paired or unpaired unique TRA/TRB clonotypes identified for 1,850-2,172 cell barcodes. As expected, individual clonotypes were characterized by evidence of germline variation, V(D)J recombination diversity, as well as somatic hypermutation and/or recombination-related mutations at V(D)J junctions (See **Figure 6** for a representative example clonotype).

**Figure 4.**
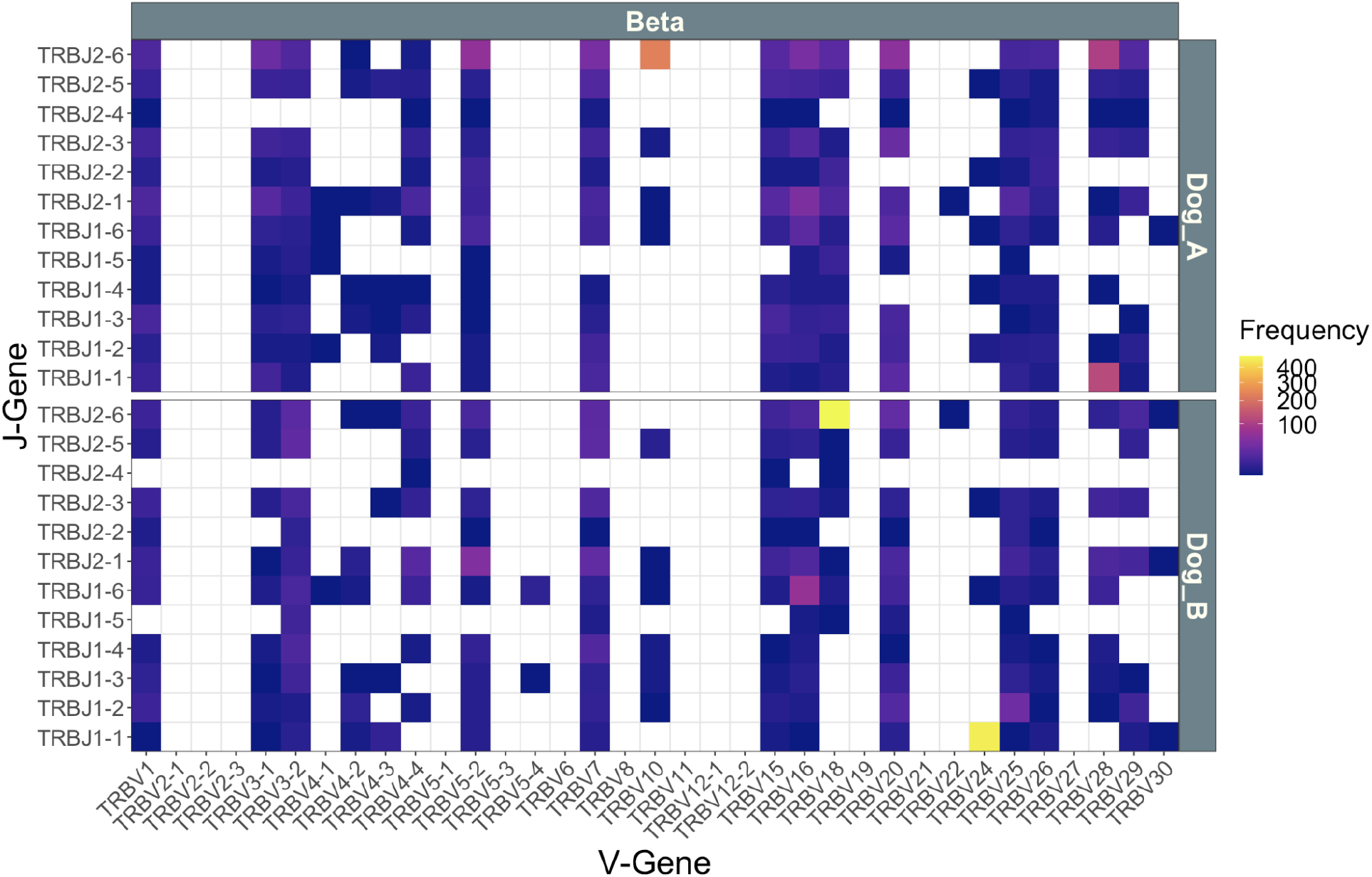
VJ Gene Segment Usage. The VJ gene combinations identified in PBMC samples for Dog_A and Dog_B are plotted along with their observed cell barcode counts for the TRB chain. The corresponding VJ gene usage for the TRA chain is shown in **Suppl Figure 3**.

**Figure 5.**
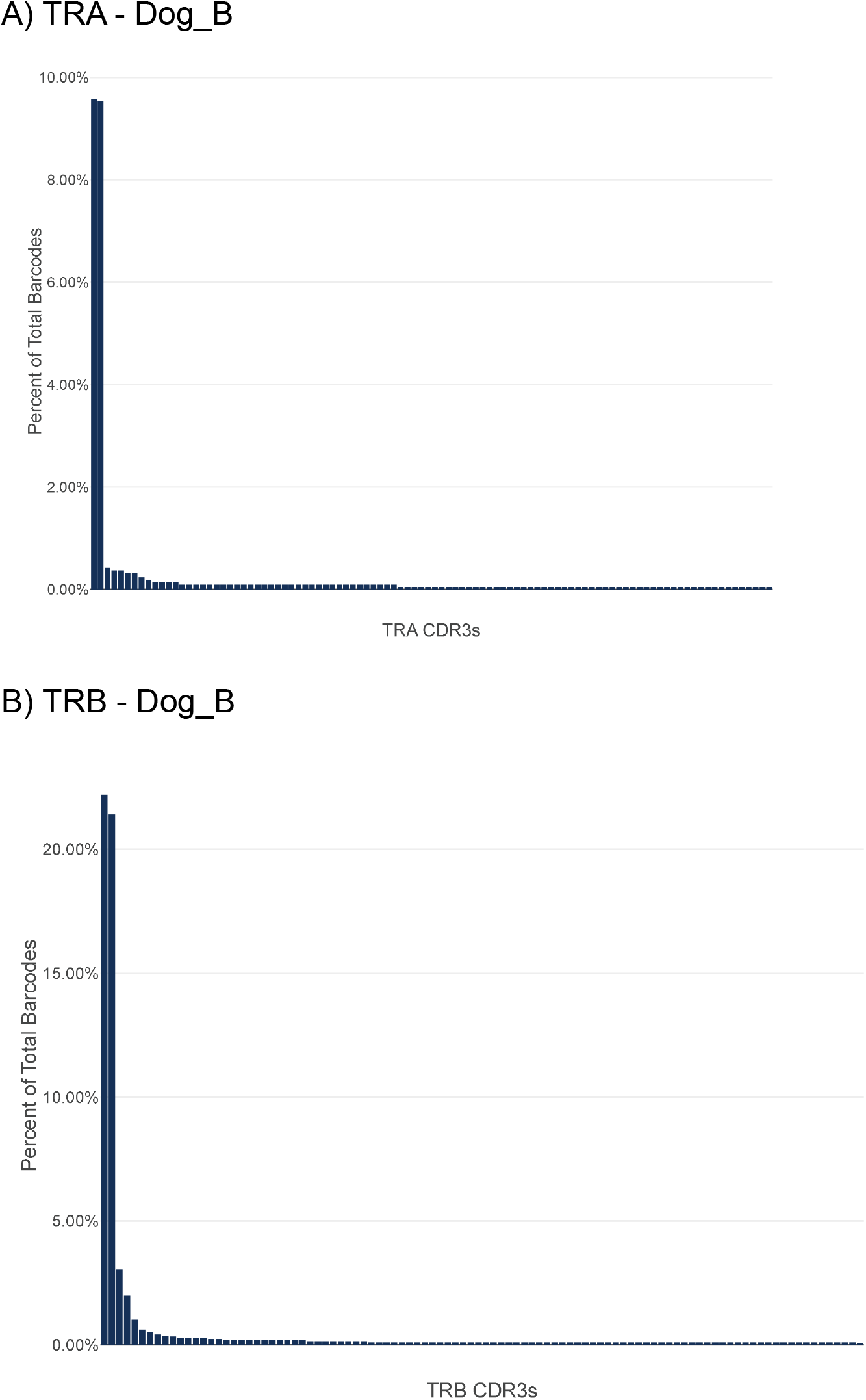
Single cell CDR3 clonotype distribution for TRA and TRB chains for Dog_B. The percent of total barcodes, for the top 100 most frequent CDR3 clonotypes, is shown for Dog_B for (A) T cell receptor alpha (TRA) and (B) T cell receptor beta (TRB). In both cases the frequency distributions are characterized by a small number of dominant clonotypes with higher frequency.

**Figure 6.**
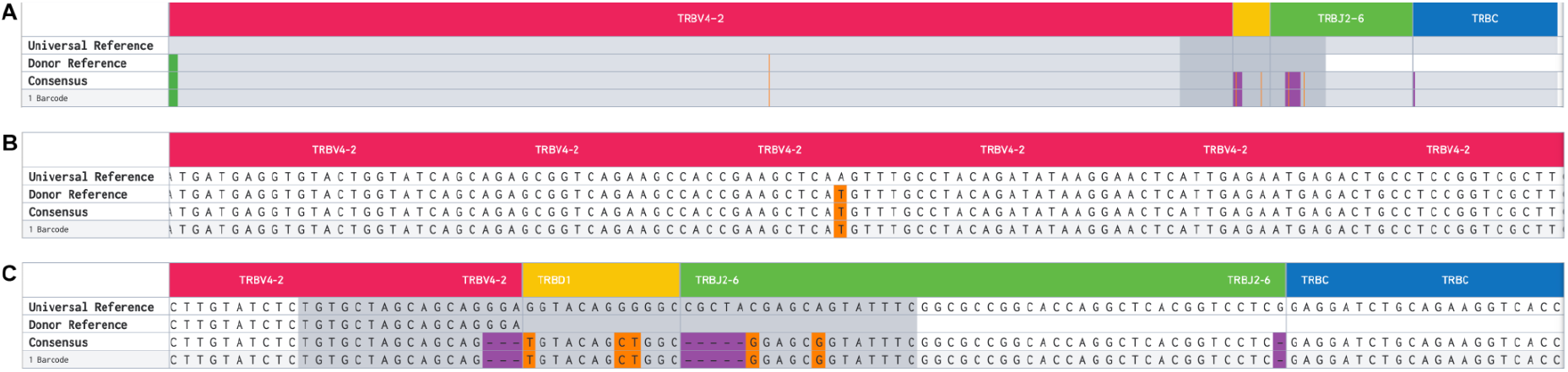
Example individual clonotype. Individual cell-specific TRA and TRB clonotypes are resolved to the nucleotide level. Illustrated here is a single such TRB clonotype from Dog_B (CDR3: CASSSVQLAERYF). (A) a specific VDJ recombination with complete TRBV4-2, TRBD1, and TRBJ2-6 and a portion of TRBC is depicted. The Universal Reference (i.e., CanFam3.1) sequence is shown in the first row. Germline variants in the analyzed sample, relative to the Universal Reference, are depicted in the Donor Reference line. The Consensus row shows additional variants from the Donor and Universal Reference that were presumably introduced during joining of gene segments. Single nucleotide changes are shown in orange and small deletions shown in purple. (B) and (C) are zoomed into the base pair level to visualize the variants.

### Integration of single cell V(D)J repertoire and gene expression data

For both dogs TSNE clustering based on gene expression patterns identified a number of distinct clusters of cells (**Figure 7**; **Suppl Figure 5**). Expression of CD3E (a general T cell marker) clearly demarcated a subset of major and minor clusters, representing a substantial fraction (∼50%) of all cells (**Figure 7A; Suppl Figure 5A**). Cell typing based on established blood cell type gene expression signatures (mapped from human genes to dog orthologs) for Dog_A identified 47.8% T cells (26.2% CD4+; 21.6% CD8+), 22.7% monocytes, 21.1% B cells, 3.5% erythroid cells, 1% dendritic cells and less than 1% of any other cell type (**Suppl Figure 5B; Suppl Table 4**). For Dog_B, cell typing identified 49.3% T cells (19.4% CD4+; 29.9% CD8+), 33.5% monocytes, 11.1% B cells, 1.2% erythroid, 1.2% NK cells and less than 1% of any other cell type (**Figure 7B; Suppl Table 4**). These cell type proportions were consistent with previously reported results in human PBMC samples^22^ and expectations for dog PBMC samples^23^ (**Figure 7; Suppl Figure 5**). Cells specifically identified with V(D)J rearrangements expressed (**Figure 7C; Suppl Figure 5C**) overlapped almost completely with those identified as CD3E-positive or as CD4/CD8 T cells. Finally, cells corresponding to the most dominant clonotypes largely overlapped with CD8 T cells.

**Figure 7.**
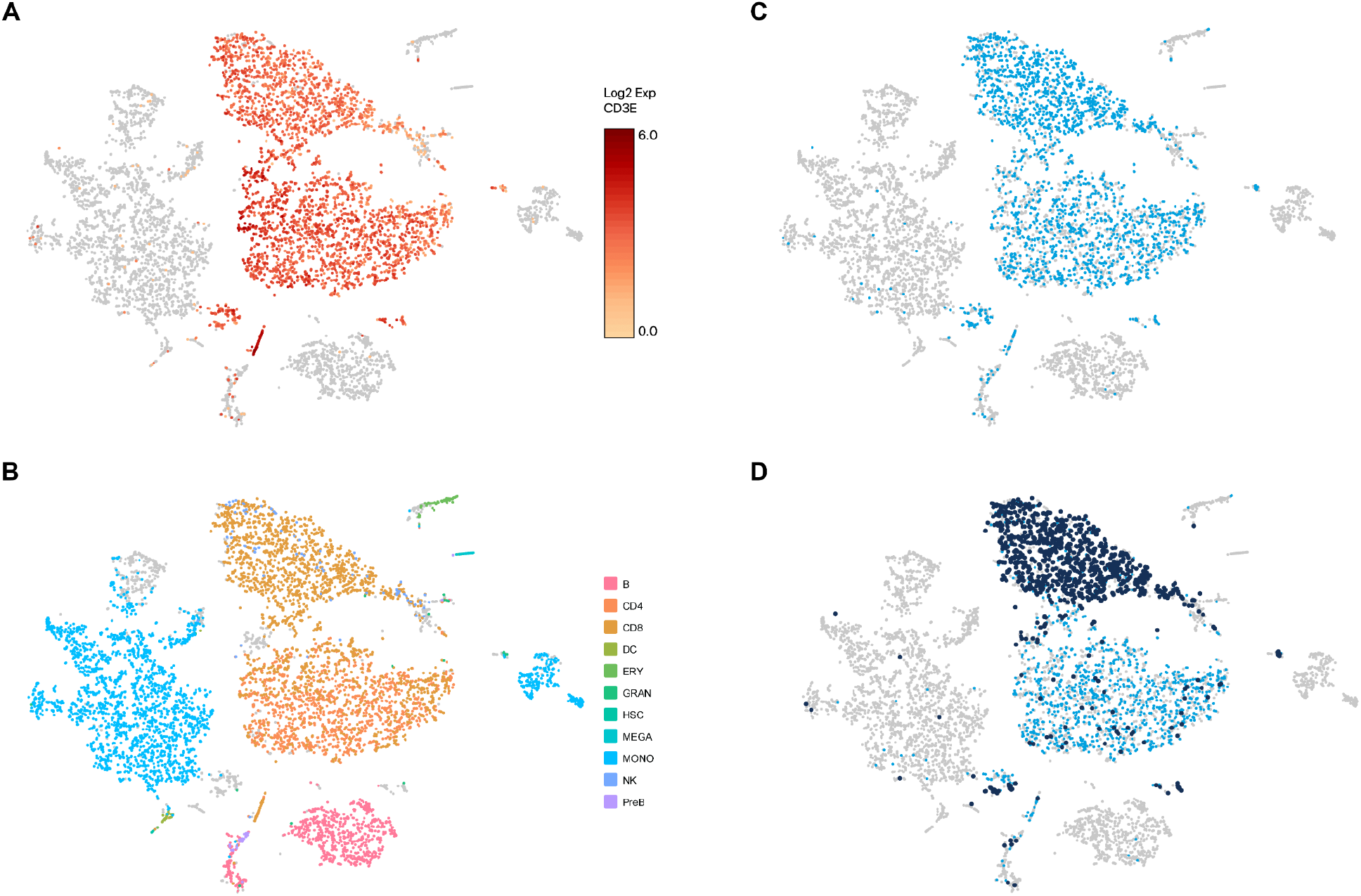
Gene expression based TSNE clustering for Dog_B categorized by CD3E expression, inferred cell type and V(D)J identification. TSNE clustering based on global gene expression patterns identified a number of distinct clusters of cells from Dog_B. (A) A T cell marker (CD3E; colored red to orange) strongly overlaps with a set of major clusters and several smaller clusters. (B) Cell types, inferred based on published expression signatures of blood cell type, identified CD4 (dark orange) and CD8 (light orange) T cell clusters largely overlapping with the CD3E-positive clusters identified in (A) as well as large monocyte (light blue), B cell (pink) clusters and smaller clusters of several other cell types. Cell types without an assignment (NA) or with less than 5 cells identified were excluded. Cells filtered based on doublet or mitochondrial filtering are shown in grey. (C) Cells identified with V(D)J rearrangements (light blue) overlap strongly with those identified as CD4/CD8 T cells or CD3E-positive. (D) Cells corresponding to the most dominant clonotypes (dark blue; **see Suppl Figure 4**) largely cluster together in the CD8 T cell cluster.

## Discussion

The biomedical community increasingly recognizes the value of companion dogs as a model system for human cancer. However, there remains a lack of the necessary reagents and methods for molecular profiling of canine clinical samples. Here we define a protocol for single cell TCR sequencing that should enhance the utility of canine samples as they relate to the study of cancer and disease. In this work we were able to successfully adapt existing protocols to perform single cell TCR profiling for two individual companion dogs. We were able to detect a majority of known V(D)J gene segments amongst our samples for both alpha and beta TCR chains. Integration with single cell transcriptome sequencing demonstrated the expected relationship between cells expressing rearranged V(D)J sequences and T cell markers or cell type inferred from global gene expression patterns. Somewhat surprisingly, both canine samples showed strong evidence of dominant clonotypes. Indeed, we observed that a number of the more frequent clonotypes could be merged based on CDR3 sequence similarity, even further emphasizing the dominant clonotype pattern (**Suppl Table 2-3**). As melanoma patients undergoing active immunotherapy treatments these patterns might indicate an active T-cell population responding to their tumors. However, such conclusions await further immune studies of these canine patient samples.

We did not detect all known V(D)J gene segments. In theory, all V, D, or J gene segments may participate in V(D)J rearrangement and form functional TCRs. However, there are a number of plausible explanations for unobserved gene segments. Missing V/J gene segments may simply be the result of insufficient sampling. D gene segments are difficult to determine due to their small size and the introduction of junction recombination mutations that make mapping to reference D gene sequences difficult. Despite the very comprehensive sequencing performed, only a few thousand cells were sequenced in relation to the very large possible diversity of T cells. This situation was compounded by the fact that the samples showed evidence of dominant clonotypes. As a result, a significant fraction of cells and sequence reads were associated with these few dominant clonotypes, decreasing our power to detect rare clonotypes. In human TCR repertoires it has been suggested that V(D)J gene segment usage is not random with some gene segments, segment combinations, and CDR3 lengths preferentially used and others rarely used^24–28^. The same could also be true in dog TCR repertoires. Finally, it is also possible that the current database of known V(D)J gene segments is incomplete. Modern canine genome assemblies^29^, future annotation work, and de novo analysis of data like ours may identify novel gene segments in the future. Profiling of additional samples will be required to further validate this approach and delineate any technological or biological gaps in V(D)J representation.

The methods developed in this study could be immediately applied to current canine immunotherapies studies to help identify predictors of response and evaluate candidate immunotherapeutic targets and efficacy of immune stimulating protocols. Future work should include adapting the protocols herein to be applicable for additional TCR loci (e.g., TRD and TRG) as well as B-cell receptor (BCR) sequencing. The ability to perform BCR sequencing would further increase the utility of canine samples, allowing for a more complete understanding of adaptive immunity in cancer, auto-immune diseases and B-cell malignancies.

## Data availability

All raw scRNAseq and scTCRseq data have been deposited at SRA:SUB9918174.

## Acknowledgements

This research was supported by the Alvin J. Siteman Cancer Center Siteman Investment Program (supported by The Foundation for Barnes-Jewish Hospital, Cancer Frontier Fund and Barnard Trust).

## Supplementary Data, Tables and Figures

**Suppl Data 1. VDJ reference sequences for use with cellranger**

**Suppl Table 1.** Detailed clinical history for dog samples.

|  | Dog_A | Dog_B |
| --- | --- | --- |
| Age | 7 | 14 |
| Sex | Female spayed | Male castrated |
| Breed | Bouvier des Flandres | American Cocker Spaniel |
| Diagnosis | Melanoma of unknown primary (superficial cervical lymph node) | Buccal mucosal melanoma |
| Treatment | Lymphadenectomy followed by deglycosylated autologous vaccine. Marginal excision of cutaneous melanoma in ipsilateral antebrachium followed by deglycosylated autologous vaccine (320 days post original diagnosis). Lymphadenectomy with lip melanoma marginal excision followed by Oncept vaccine (1698 days post original diagnosis). | Marginal excision of primary lesion. Marginal excision of recurrent now amelanotic melanoma with lymphadenectomy followed by deglycosylated autologous vaccine (95 days post original diagnosis). Marginal excision of recurrent amelanotic melanoma followed by deglycosylated autologous vaccine (598 days post original diagnosis). |
| Concurrent Disease(s) | Bilateral glaucoma | Protein losing nephropathy, sebaceous adenomas |
| Cause of Death | Metastatic melanoma (multifocal skin, heart, kidney, lung, liver, inguinal lymph node) | Hemoabdomen (unknown cause). Pulmonary metastasis and urogenital transitional cell carcinoma on necropsy but determined to not be the cause of death. |
| Survival (days) | 1858 | 741 |

**Suppl Table 2. TCR clonotypes observed for Dog_A**

**Suppl Table 3. TCR clonotypes observed for Dog_B**

**Suppl Table 4.** Cell typing results.

|  |  | **Cell counts (%)** |  |
| --- | --- | --- | --- |
| **Cell Type** | **Cell Abbrev.** | **Dog_A** | **Dog_B** |
| B-Cell | B | 1323 (21.1) | 746 (11.1) |
| Basophil | BASO | 10 (0.2) | 1 (0.01) |
| CD4+ T-cell | CD4 | 1639 (26.2) | 1296 (19.4) |
| CD8+ T-cell | CD8 | 1350 (21.6) | 2000 (29.9) |
| Common Myeloid Progenitor | CMP | 2 (0.03) | 2 (0.03) |
| Dendritic cell | DC | 63 (1.0) | 36 (0.5) |
| Eosinophil | EOS | 3 (0.05) | 0 (0.0) |
| Erythroid | ERY | 217 (3.5) | 81 (1.2) |
| Granulocyte | GRAN | 11 (0.2) | 25 (0.4) |
| Hematopoietic Stem Cell | HSC | 2 (0.03) | 6 (0.09) |
| Megakaryocyte | MEGA | 14 (0.22) | 28 (0.42) |
| Megakaryocyte/Erythroid Progenitor | MEP | 2 (0.03) | 2 (0.03) |
| Monocyte | MONO | 1421 (22.7) | 2240 (33.5) |
| Not Available/Not Determined | NA | 140 (2.2) | 115 (1.7) |
| Natural Killer | NK | 20 (0.3) | 78 (1.2) |
| Precursor B-Cell | PreB | 44 (0.7) | 38 (0.6) |
| Totals (filtered) | 16 cell types | 6291 | 6694 |

**Supplementary Figure 1.**
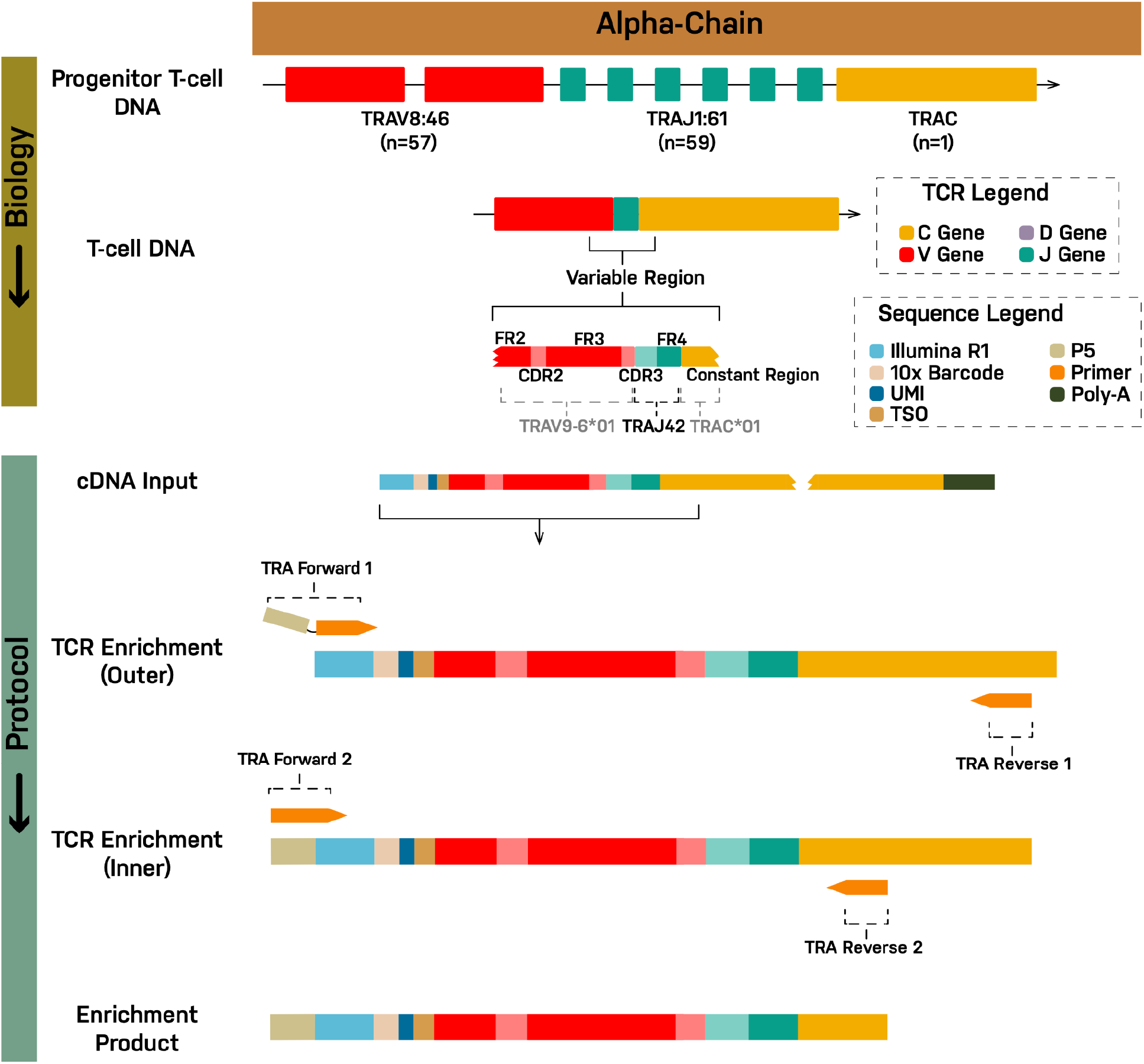
The TCR V(D)J enrichment strategy is depicted for the alpha chain (See **Figure 1** for beta chain). At top, the genomic un-rearranged TRA locus is shown. During T cell development from progenitor T cells to mature T cells, individual V, J and C gene segments are rearranged by somatic recombination to produce a functional TRA locus. Transcription and splicing produce a pre-mRNA and then mRNA for the complete VJC transcript sequence (not shown). In the modified 10X protocol, mRNA (including TRA mRNA) is converted to cDNA. TRA cDNA is then amplified using a nested PCR design. The two forward primers from the 10x protocol were left unchanged. In the first cycle, the forward primer (TRA Forward 1) primes off the Illumina read 1 (R1) sequencing adapter that is incorporated during generation of cDNA. TRA Forward 1 includes a tail sequence consisting of the P5 priming site used in Illumina sequencing. In the second cycle, the forward primer (TRA Forward 2) primes off the P5 sequence and a portion of R1. The first reverse primer (TRA Reverse 1, Outer) primes off the constant (C) region gene segment. The second reverse primer (TRA Reverse 2, Inner) similarly primes off the C region at an inner, 5’ position relative to the outer primer. The alpha chain primer design was based off of a dog TCR alpha rearranged partial mRNA (GenBank: M97511.1) which was extended to include (from 3’ to 5’) the R1 adaptor, 10x cell barcode, UMI, TSO, V, J, and C gene segments. The constructed cDNA sequence was then used as input to primer3plus (4.0), with forward primers provided as described above, and a target region for reverse primer specified in the C region. The product of the first (outer) design was used as input for the second (inner) design.

**Supplementary Figure 2.**
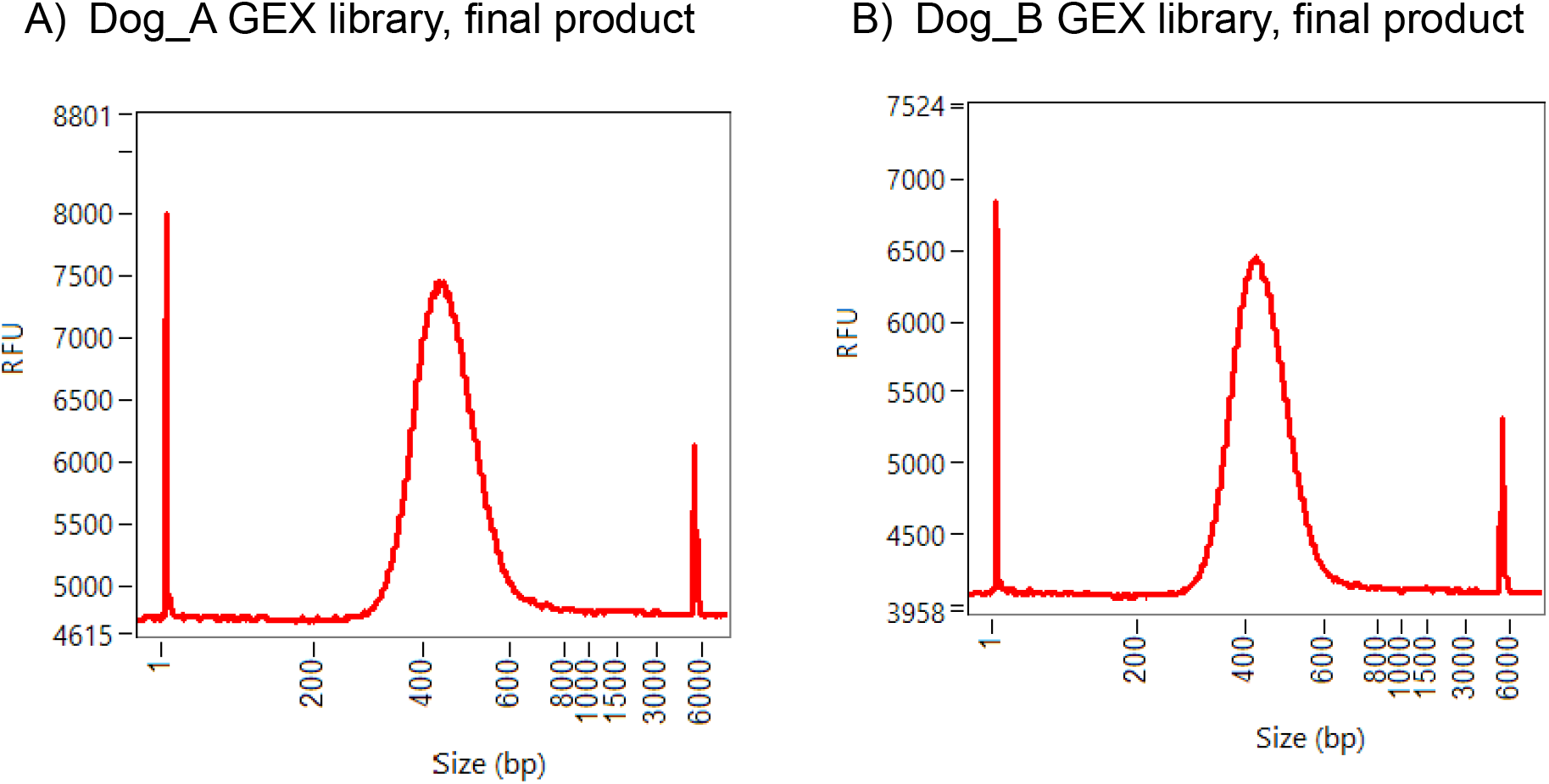
Analytic traces of the expression libraries for Dog_A and Dog_B by the Fragment Analyzer. The final purified 5’ gene expression(GEX) libraries were analyzed on Agilent Fragment Analyzer using the NGS Fragment kit (DNF-473-500). The traces show normal gene expression libraries for (A) Dog_A and (B) Dog_B.

**Supplementary Figure 3.**
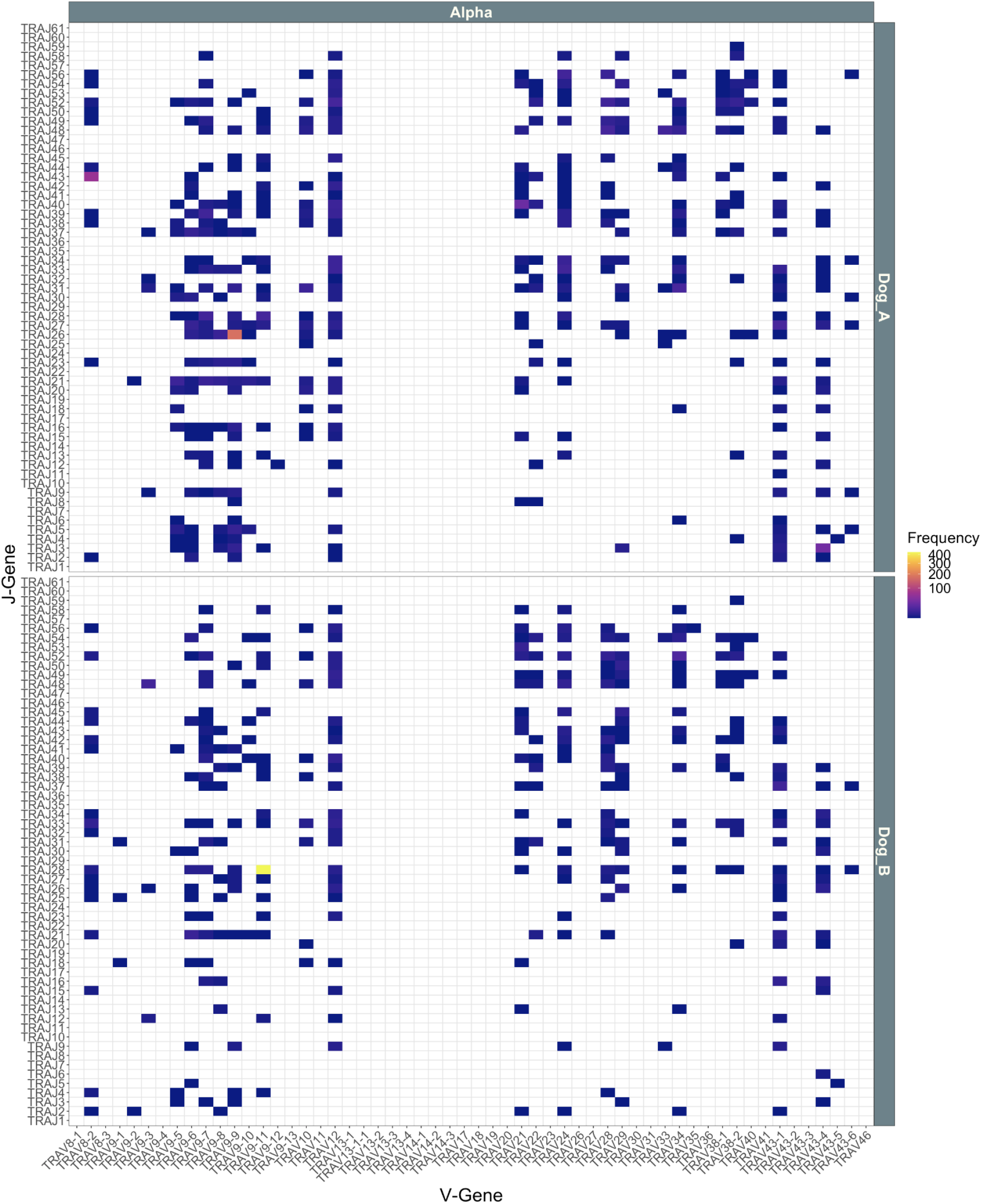
V(D)J Gene Segment Usage – alpha. The VJ gene combinations identified in PBMC samples for Dog_A and Dog_B are plotted along with their observed cell barcode counts for the TRA chain.

**Supplementary Figure 4.**
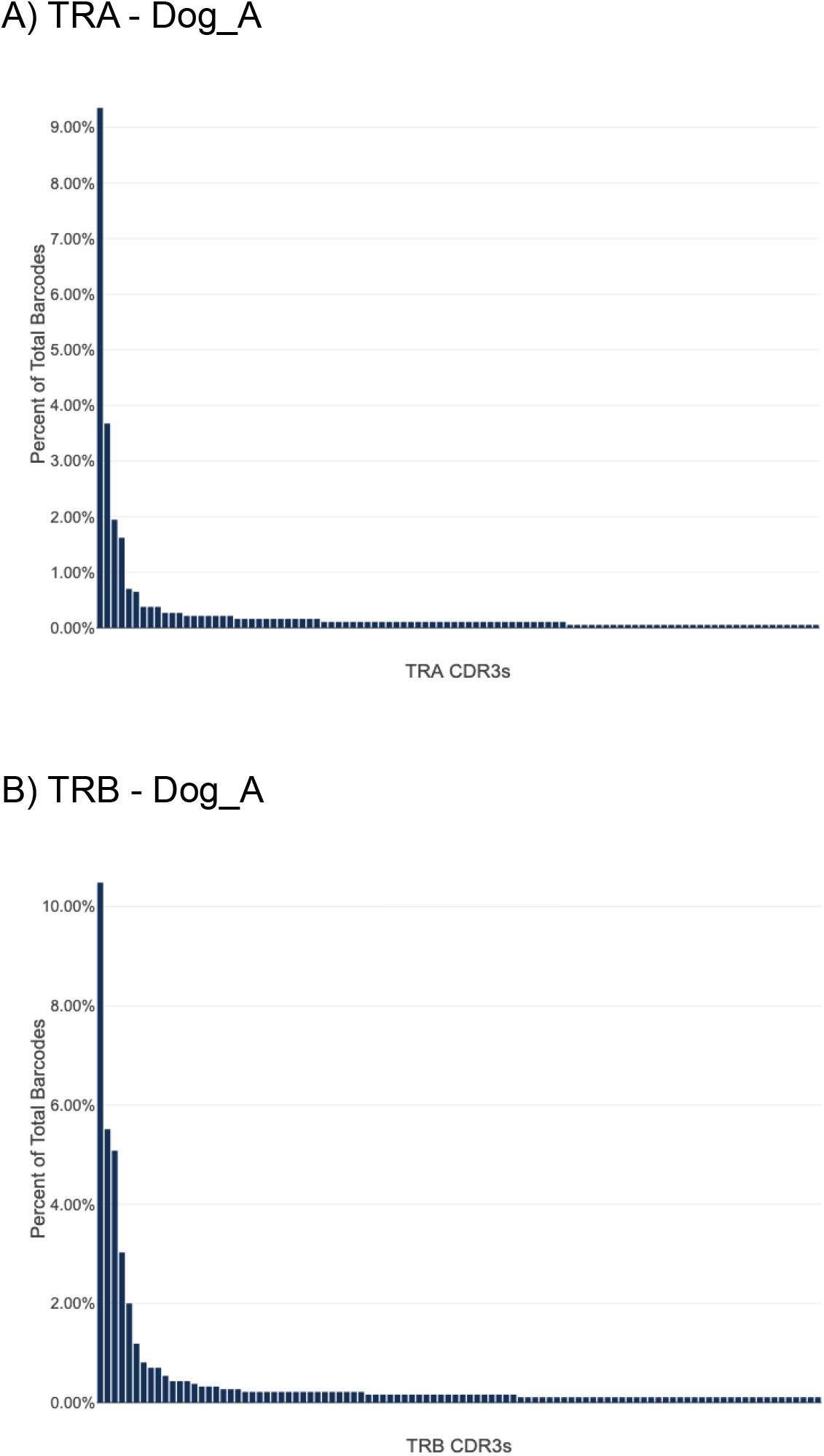
Single cell CDR3 clonotype distribution for TRA and TRB chains for Dog_A. The percent of total barcodes, for the top 100 most frequent CDR3 clonotypes, is depicted for Dog_A for (A) T cell receptor alpha (TRA) and (B) T cell receptor beta (TRB). In both cases the frequency distributions are characterized by a small number of dominant clonotypes with higher frequency.

**Supplementary Figure 5.**
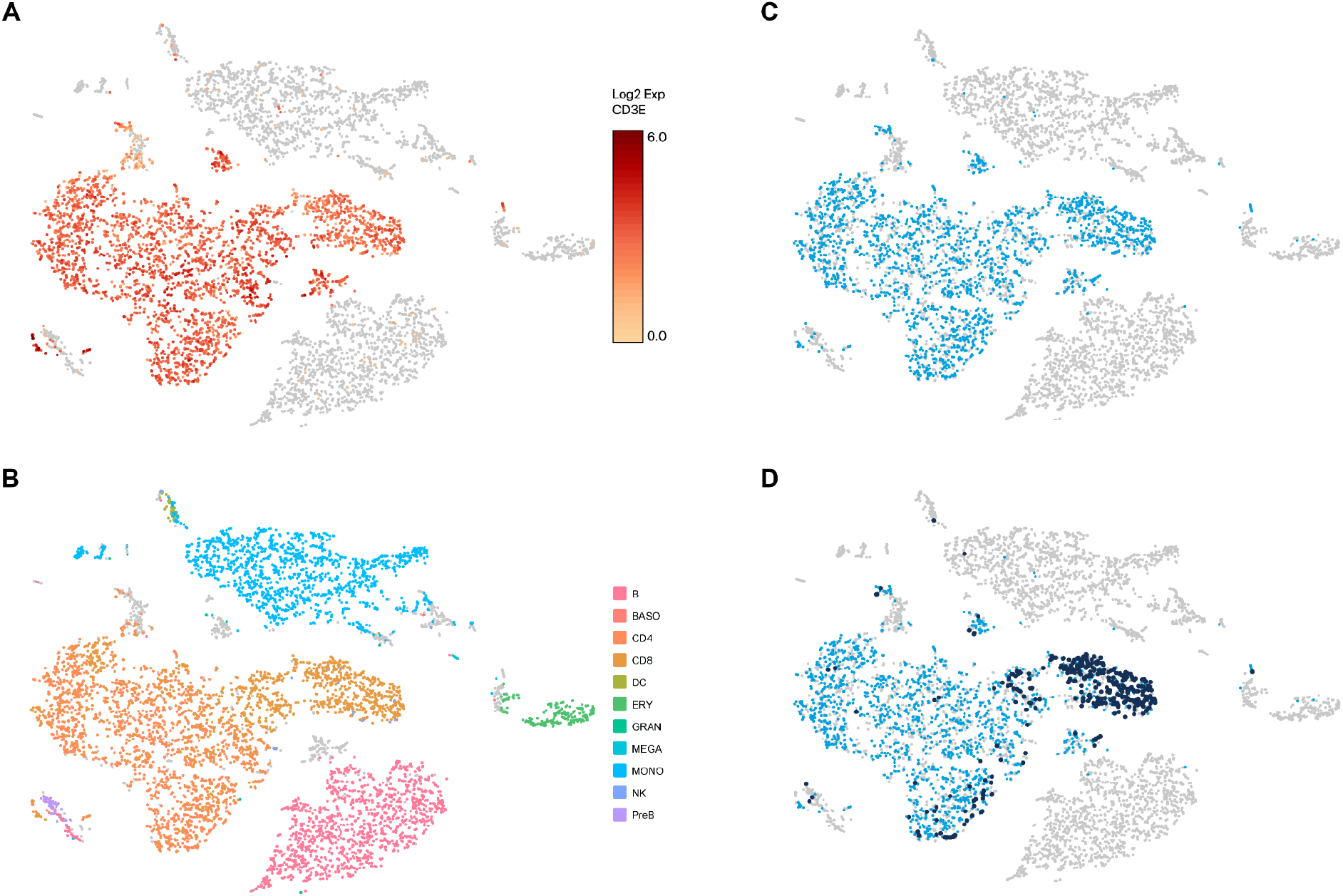
Gene expression based TSNE clustering for Dog_A categorized by CD3E expression, inferred cell type and V(D)J identification. TSNE clustering based on global gene expression patterns identified a number of distinct clusters of cells from Dog_A. (A) A T cell marker (CD3E; colored red to orange) strongly overlaps with a set of major clusters and several smaller clusters. (B) Cell types, inferred based on published expression signatures of blood cell type, identified CD4 (dark orange) and CD8 (light orange) T cell clusters largely overlapping with the CD3E-positive clusters identified in (A) as well as large monocyte (light blue), B cell (pink) clusters and smaller clusters of several other cell types. Cell types without an assignment (NA) or with less than 5 cells identified were excluded. Cells filtered based on doublet or mitochondrial filtering are shown in grey. (C) Cells identified with V(D)J rearrangements (light blue) overlap strongly with those identified as CD4/CD8 T cells or CD3E-positive. (D) Cells corresponding to the most dominant clonotypes (dark blue; **see Suppl Figure 4**) largely cluster together in the CD8 T cell cluster.

